# BRWD3 promotes KDM5 degradation to maintain H3K4 methylation levels

**DOI:** 10.1101/2023.03.28.534572

**Authors:** Dongsheng Han, Samantha H. Schaffner, Jonathan P. Davies, Mary Lauren Benton, Lars Plate, Jared T. Nordman

**Affiliations:** Department of Biological Sciences, Vanderbilt University, Nashville, TN, 37212, USA; Department of Chemistry, Vanderbilt University, Nashville, TN, 37212, USA; Department of Computer Science, Baylor University, Waco, TX 76798, USA

## Abstract

Histone modifications are critical for regulating chromatin structure and gene expression. Dysregulation of histone modifications likely contributes to disease states and cancer. Depletion of the chromatin-binding protein BRWD3, a known substrate-specificity factor of the Cul4-DDB1 E3 ubiquitin ligase complex, results in increased in H3K4me1 levels. The underlying mechanism linking BRWD3 and H3K4 methylation, however, has yet to be defined. Here, we show that depleting BRWD3 not only causes an increase in H3K4me1 levels, but also causes a decrease in H3K4me3 levels, indicating that BRWD3 influences H3K4 methylation more broadly. Using immunoprecipitation coupled to quantitative mass spectrometry, we identified an interaction between BRWD3 and the H3K4-specific demethylase 5 (KDM5/Lid), an enzyme that removes tri- and di- methyl marks from H3K4. Moreover, analysis of ChIP-seq data revealed that BRWD3 and KDM5 are significantly co- localized throughout the genome and that sites of H3K4me3 are highly enriched at BRWD3 binding sites. We show that BRWD3 promotes K48-linked polyubiquitination and degradation of KDM5 and that KDM5 degradation is dependent on both BRWD3 and Cul4. Critically, depleting KDM5 fully restores altered H3K4me3 levels and partially restores H3K4me1 levels upon BRWD3 depletion. Together, our results demonstrate that BRWD3 regulates KDM5 activity to balance H3K4 methylation levels.

## Introduction

Epigenetic regulation of gene expression through histone modifications can alter chromatin structure and function without any alteration in DNA sequence. This allows for changes in gene expression that can ultimately drive cell differentiation and disease states (1, 2). The N-terminal tails of histones undergo numerous modifications including methylation, acetylation, and phosphorylation, among others (3). These modifications affect chromatin structure and serve as binding platforms for transcription factors, chromatin remodelers, and histone modification modifying enzymes (3).

One histone modification that is commonly associated with actively transcribed genes is histone H3 lysine 4 methylation (H3K4me) (1, 4, 5). H3K4 can be methylated up to three times, resulting in mono, di-, or tri-methylation (H3K4me1, H3K4me2 or H3K4me3, respectively) (6). While H3K4me3 is primarily enriched at gene promoters and around transcription start sites, H3K4me2 is spread throughout intragenic and intergenic regions while H3K4me1 is commonly found at enhancer regions (7–9). Normal H3K4 methylation patterns are important for development and embryogenesis and the steady state of H3K4 methylation is controlled by a balance between H3K4-specific methyltransferases (KMTs) that add methyl groups and H3K4-specific lysine demethylases (KDMs) that remove methyl groups (6, 10). Set1-domain containing complexes known as COMPASS (Complex of Proteins Associated with Set1), catalyzes mono-, di, and trimethylation of histone H3K4 (6). On the other hand, there are two families of demethylases, KDM1 and KDM5 that remove methyl groups form H3K4. While the KDM1 family of enzymes removes methyl groups from H3K4me1 and H3K4me2, KDM5 family preferentially removes the methyl group from H3K4me3 and to a lesser extent H3K4me2 (11–14). The enzymes that control H3K4 methylation are essential for normal cellular function (4). For example, altered expression of human *KDM5* genes have been implicated in various cancers and *KDM5* mutations are linked to intellectual disorders (15–18). Although the enzymes control H3K4 methylation are well defined, how these enzymes are regulated to balance H3K4 methylation status is largely unknown.

The evolutionary conserved protein, Bromodomain and WD repeat-containing protein 3 (BRWD3), binds directly to H3K4 methylation *in vitro* through a cryptic Tudor domain (19). The BRWD3 homolog also has two bromodomains that bind to acetylated histones with simultaneous binding of H3K4 methylation (20). Interestingly, loss of BRWD3 function causes an increase in H3K4me1 levels in Drosophila, but the underlying mechanism is completely unknown (19) BRWD3 functions as a substrate receptor of the Cul4 E3 ubiquitin ligase complex (21–23). While Cul4 is known to target many DNA replication and cell cycle proteins for degradation, how BRWD3 contributes to epigenetic maintenance and if ubiquitin targeting activity is involved is not clear (21, 24, 25). Mutations in the *BRWD3* family of genes were recently identified as the cause of a neurodevelopmental disorder and altered *BRWD3* expression has been found in various cancers (26–30). Thus, defining the molecular function of BRWD3 in epigenetic maintenance could be an important first step for understanding the molecular causes of disease states and identifying potential therapeutic targets.

Drosophila is an ideal system to understand how BRWD3 regulates H3K4 methylation (31). Drosophila has a single BRWD3 homolog, whereas humans have three BRWD3 orthologs (BRWD1, BRWD2, and BRWD3). Furthermore, H3K4 methylation is streamlined in Drosophila relative to humans. Drosophila has three H3K4 methyltransferase complexes (Set1/COMPASS, Tri, and Trr), whereas humans have six (SET1A, SET1B, MLL1, MLL2, MLL3, and MLL4) (6). For H3K4me3/me2 demethylation, Drosophila has a single KDM5 family member (KDM5/Lid), whereas humans have up to four KDM5 orthologs (KDM5A, KDM5B, KDM5C, and KDM5D) (16).

In this study, we show BRWD3 has a global effect on H3K4 methylation levels. While loss of BRWD3 function has been shown to promote an increase in H3K4me1 levels, we find that H3K4me3 levels are reduced upon depletion of BRWD3. Strikingly, we identify KDM5 as a BRWD3- associated protein. *In vivo* ubiquitination assays reveal that BRWD3 promotes K48-linked polyubiquitination on KDM5. Furthermore, KDM5 is rapidly degraded in a BRWD3- and Cul4- dependent manner. Critically, co-depletion of KDM5 can restore H3K4me3 levels upon BRWD3 depletion. We also find that *BRWD3* is a *su(var)* gene and loss of single copy of *KDM5* can suppress the *BRWD3* Su(var) phenotype. Taken together, our work supports a model in which BRWD3 controls the stability of KDM5 to regulate demethylation of H3K4me3 and maintain overall H3K4 methylation levels.

## Results

### BRWD3 affects H3K4 mono, di and tri methylation levels

In Drosophila, BRWD3 is required to maintain H3K4me1 levels although the mechanism is unknown (19). Due to the dynamic regulation of H3K4me1, me2 and me3, we were curious whether BRWD3 also influenced H3K4me3 and/or me2 methylation levels. In addition, we wanted to determine if BRWD3 control of H3K4 methylation is dependent on Cul4 E3 ubiquitin ligase given BRWD3 is a substrate specificity factor for the Cul4 ubiquitin ligase complex (21). To measure the relative level of H3K4 methylation marks, we depleted BRWD3 and Cul4 in Drosophila S2 cells and measured H3K4me1, me2 and me3 levels by quantitative immunofluorescence (IF). Consistent with previous results, BRWD3 depletion caused an increased H3K4me1 levels relative to negative control cells (GFP) (Fig. 1A). We found that BRWD3 depletion caused a decrease in H3K4me3 levels (Fig 1B) and an increase in H3K4me2 levels (Fig. S1). Importantly, Cul4 depletion resulted in similar changes in H3K4me1 (Fig. 1A), me2 (Fig. S1A), and me3 levels (Fig. 1B). Taken together, these results demonstrate that BRWD3 affects pan H3K4 methylation levels and likely does so in a ubiquitination-dependent manner.

**Figure 1.**
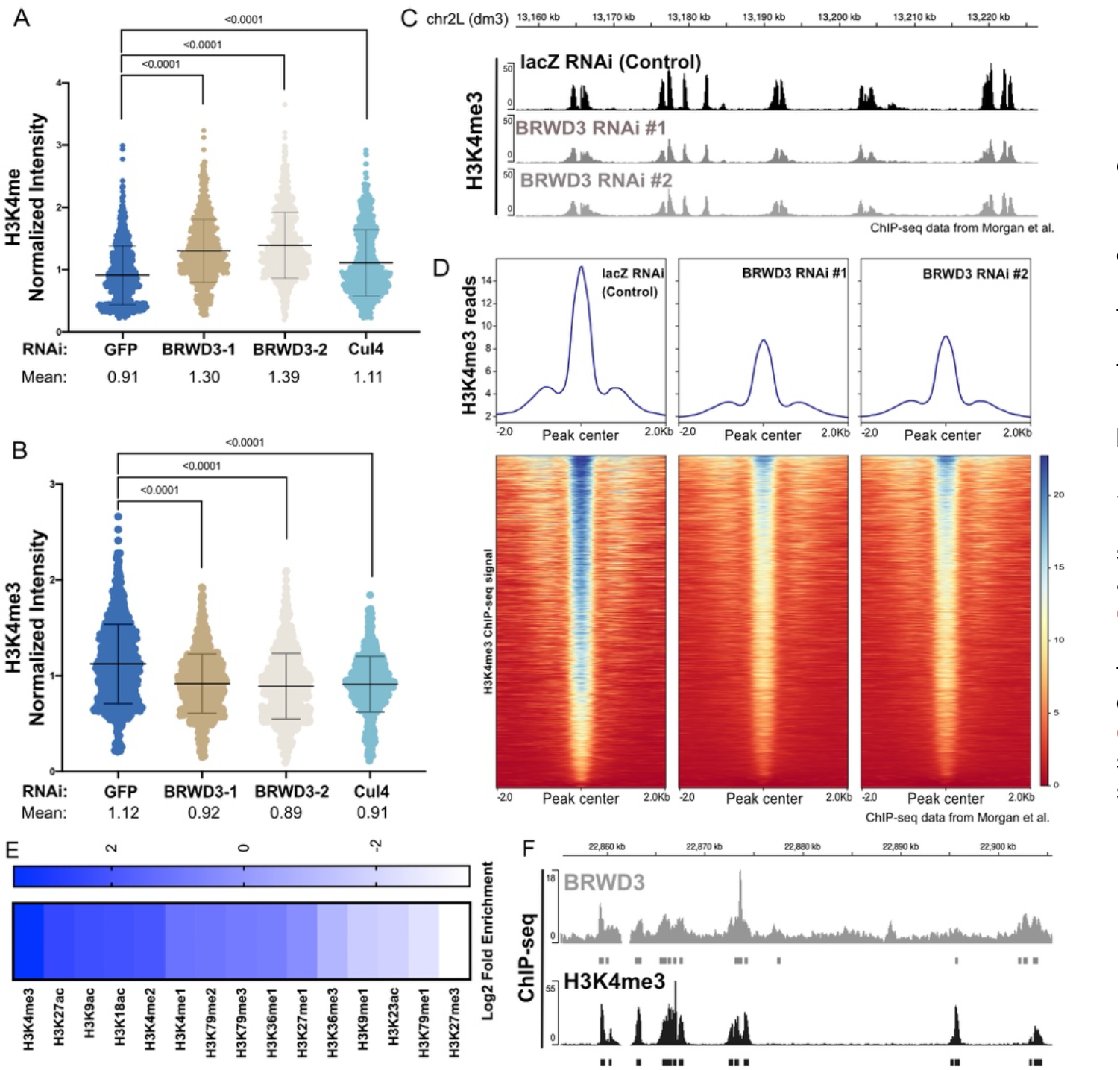
BRWD3 affects H3K4 methylation status. Quantification of H3K4me1 **(A)** and H3K4me3 **(B)** intensity using quantitative IF in Drosophila S2 cells. Each dot presents the H3K4 methylation intensity per nucleus normalized to the total DNA content. Each distribution represents the signal intensities of 1000 randomly selected cells from three biological replicates. p < 0.0001 using an ANOVA one-way analysis with Tukey’s multiple comparisons test. **(C)** Representative H3K4me3 ChIP-seq profiles from negative control (lacZ) or BRWD3 depletion using two dsRNAs in S2 cells. Data was from (19, 32). **(D)** Enrichment heatmap of H3K4me3 ChIP- seq signal sorted by mean occupancy around the centers of all H3K4me3 peaks. **(E)** Enrichment of H3 marks within BRWD3 binding sites. Log2 enrichment for observed overlap relative to expected overlap for each mark peak set is shown. **(F)** Representative view showing the similarity of BRWD3 and H3K4me3 ChIP- seq tracks. Data was from (19).

To test further if BRWD3 regulates H3K4me3 levels using an independent approach, we re-analyzed previously published H3K4me3 ChIP-seq data generated in control (lacZ) and BRWD3-depleted Drosophila S2 cells (19). Consistent with our IF data, depletion of BRWD3 caused a reduction in H3K4me3 ChIP-seq reads at H3K4me3 peak regions relative to negative control cells and the reduction of H3K4me3 signal was ubiquitous throughout the genome (Fig. 1C). To quantify the change in H3K4me3 ChIP-seq read counts, we centered all H3K4me3 peaks and quantified the read counts for each peak (Fig. 1D). We found that BRWD3 depletion caused a ∼50% reduction of H3K4me3 ChIP-seq read counts. Thus, together with our IF results, we conclude that BRWD3 regulates not only H3K4me1, but also me2 and me3 levels.

BRWD3 homologs have a cryptic tudor domain that is critical for binding to methylated H3 and two bromo domains necessary for binding to acetylated H3 (19). The human BRWD2 homolog PHIP binds to both H3K4 methylated histones and acetylated histone H3 peptides *in vitro* (19, 20). Given that BRWD3 affects H3K4me, me2, and me3 levels in vivo, we wanted to determine if BRWD3 associates with chromatin containing these specific histone modifications throughout the genome. To this end, we evaluated the overlap between BRWD3 and all available histone modifications available through the modENCODE consortium (33, 34). For each dataset, we compared the observed overlap with the expected overlap obtained from 1,000 randomly shuffled sets of peaks (35, 36). This allowed us to calculate both the enrichment and significance of any overlap. Overall, active histone modifications were significantly enriched around BRWD3 binding sites, while repressive histone marks were significantly depleted, consistent with previous observations that BRWD3 is associated with active histone marks (Fig. 1E) (19). Interestingly, H3K4me3 was the most significantly enriched mark at BRWD3 binding sites among all tested H3 modifications (p value = 0.001, log2-fold change = 3.48) (Fig. 1E, 1F). Importantly, H3K4me3 was enriched at a higher level than H3K4me2 and H3K4me1, which mirrors the *in vitro* binding affinity of the BRWD3 cryptic tudor domain for these marks (Fig. 1E) (19, 20). Additionally, H3K27ac, H3K9ac and H3K18ac were also enriched in BRWD3 binding sites, consistent with the BRWD3 bromodomain binding to these modifications in combination with H3K4 methylation *in vitro* (20). Together, these data suggest that, while BRWD3 associates with methylated H3K4 *in vitro*, BRWD3 preferentially associates with H3K4me3 in *vivo*.

### BRWD3 associates with the H3K4me3 histone demethylase KDM5/Lid

H3K4 methylation levels are controlled by both histone methyltransferase and histone demethylase activities(6). To explore the mechanism by which BRWD3 affects H3K4 methylation status, we used immunoprecipitation followed by quantitative mass spectrometry (IP-qMS) to identify BRWD3 interacting proteins that could potentially regulate H3K4 methylation levels. To this end, we constructed an endogenously HA-tagged BRWD3 fly line using the CRISPR- Cas9-based knock-in method (37, 38). Loss of BRWD3 function is lethal in Drosophila (39, 40). The endogenously tagged *BRWD3-HA* homozygous line, however, produced a protein of the expected size and was viable indicating that BRWD-HA is functional (Fig. S2). We immunoprecipitated BRWD3-HA from benzonase-digested embryo extracts, labeled IP material with tandem mass tags (TMT) and identified BRWD3-interacting proteins by tandem qMS (Fig. 2A and Table S1). We observed a significant enrichment of Cul4, Pic/DDB1, Roc1A that are known components of the Cul4 E3 ubiquitin ligase complex (Fig. 2B) (23). Interestingly, we also identified Nedd8, which covalently modifies Cul4 to active Cul4 ubiquitin ligase activity, indicating BRWD3 can associate with activated Cul4 (Fig. 2B) (41). Strikingly, we identified the H3K4-specific lysine demethylase KDM5/Lid as a BRWD3-interacting protein (Fig. 2B).

**Figure 2.**
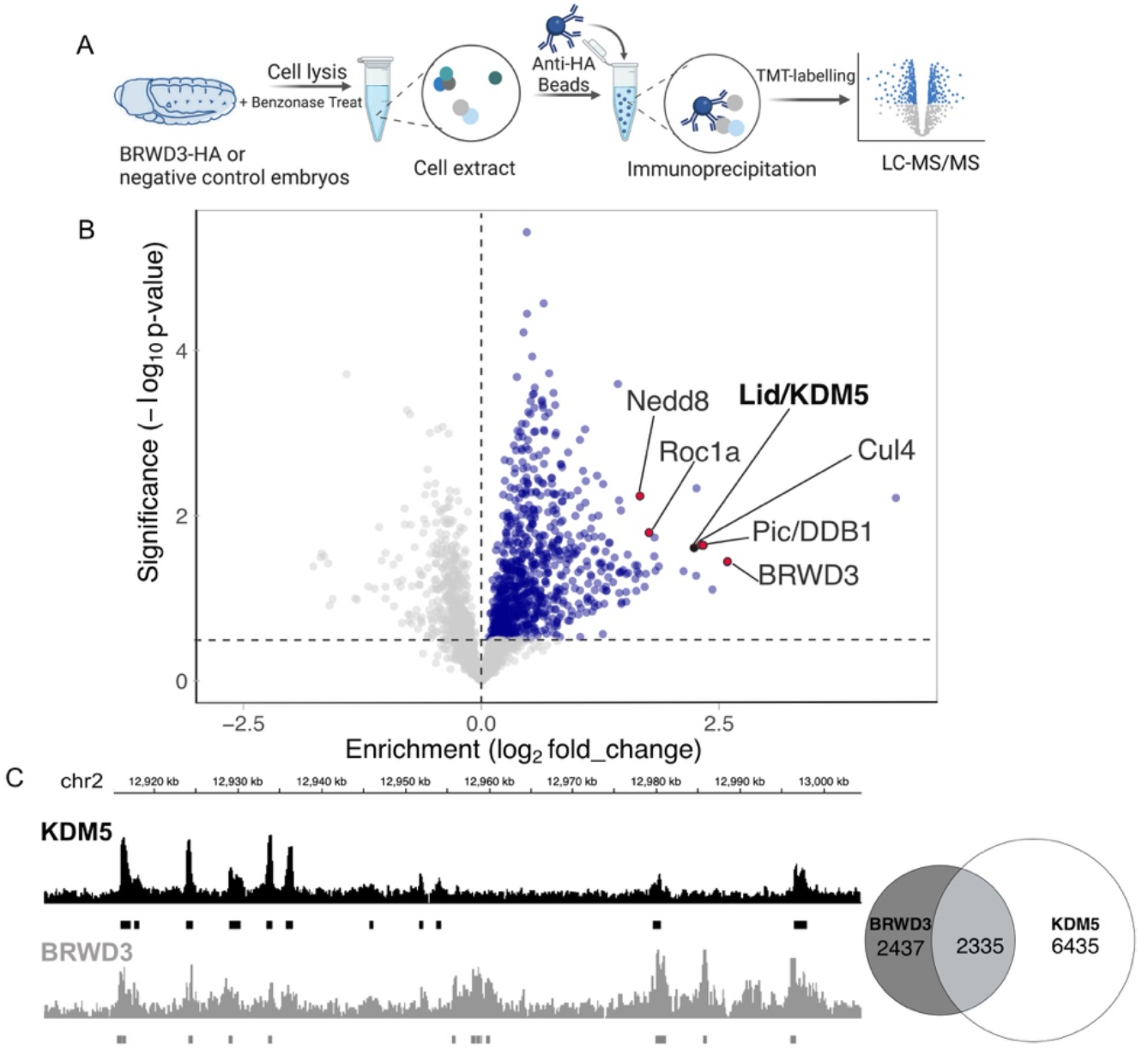
BRWD3 and KDM5 are in the same complex and bind the same genomic regions. **(A)** Workflow of BRWD3-HA immunoprecipitations from Drosophila embryos followed by mass spectrometry. **(B)** Volcano plot of the BRWD3-IP-TMT-MS results. Blue dots represent hits significantly enriched in BRWD3-HA-IP compared to the negative control. The red dots represent the Cul4-DDB1-BRWD3 E3 ubiquitin ligase complex. **(C)** Representative view showing the similarity of BRWD3 and KDM5 ChIP-seq tracks (left). Venn plot quantifying the overlap between BRWD3 and KDM5 binding sites on the genome, p<0.0001 (right).

Given the association between BRWD3 and KDM5, we next asked if BRWD3 and KDM5 share similar binding sites on chromatin. To this end, we analyzed previously published ChIP-seq data for BRWD3 (from Drosophila S2 cells) and KDM5 (from Drosophila adults) (19, 32). Even though these data sets are from desperate developmental samples, we found that ∼50% of BRWD3 binding sites overlap with KDM5 sites (Fig. 2C). Taken together, we conclude that BRWD3 binding sites show significant overlap with KDM5 sites genome wide. Given that KDM5 can sequentially remove methyl group from H3K4me3 to convert it to H3K4me2 and further to H3K4me1, we hypothesize that BRWD3 influences H3K4me3 levels by regulating KDM5 activity (13, 42).

### BRWD3 promotes KDM5 ubiquitination and degradation

Our results show that BRWD3 associates with KDM5 and is required for normal H3K4me3 levels. Given that BRWD3 is a substrate specificity factor for the Cul4 E3 ubiquitin ligase, BRWD3 could control H3K4me3 levels by targeting KDM5 for ubiquitination and possibly degradation. Thus, upon depletion of BRWD3, KDM5 levels may become elevated resulting in excessive H3K4me3 demethylase activity. To test this hypothesis, we first assessed whether KDM5 ubiquitination was dependent on BRWD3 *in vivo* (see Methods). In these experiments, we co-expressed HA-tagged KDM5 with FLAG-ubiquitin in Drosophila S2 cells and analyzed KDM5 ubiquitination status following a HA-IP under denaturing conditions. We detected a low level of ubiquitination of KDM5 in control cells (Fig. 3A). BRWD3 depletion, however, did not cause a significant effect on KDM5 ubiquitination, which would be expected if multiple ubiquitin ligases targeted KDM5 (Fig. 3A). In contrast, overexpression of BRWD3 greatly enhanced KDM5 ubiquitination (Fig.3A). By using ubiquitin-linkage-specific antibodies we found that BRWD3 promotes K48-linked polyubiquitination of KDM5 (Fig. 3B). We conclude that BRWD3 promotes K48- linked ubiquitination of KDM5, suggesting that BRWD3 could affect KDM5 stability.

**Figure 3.**
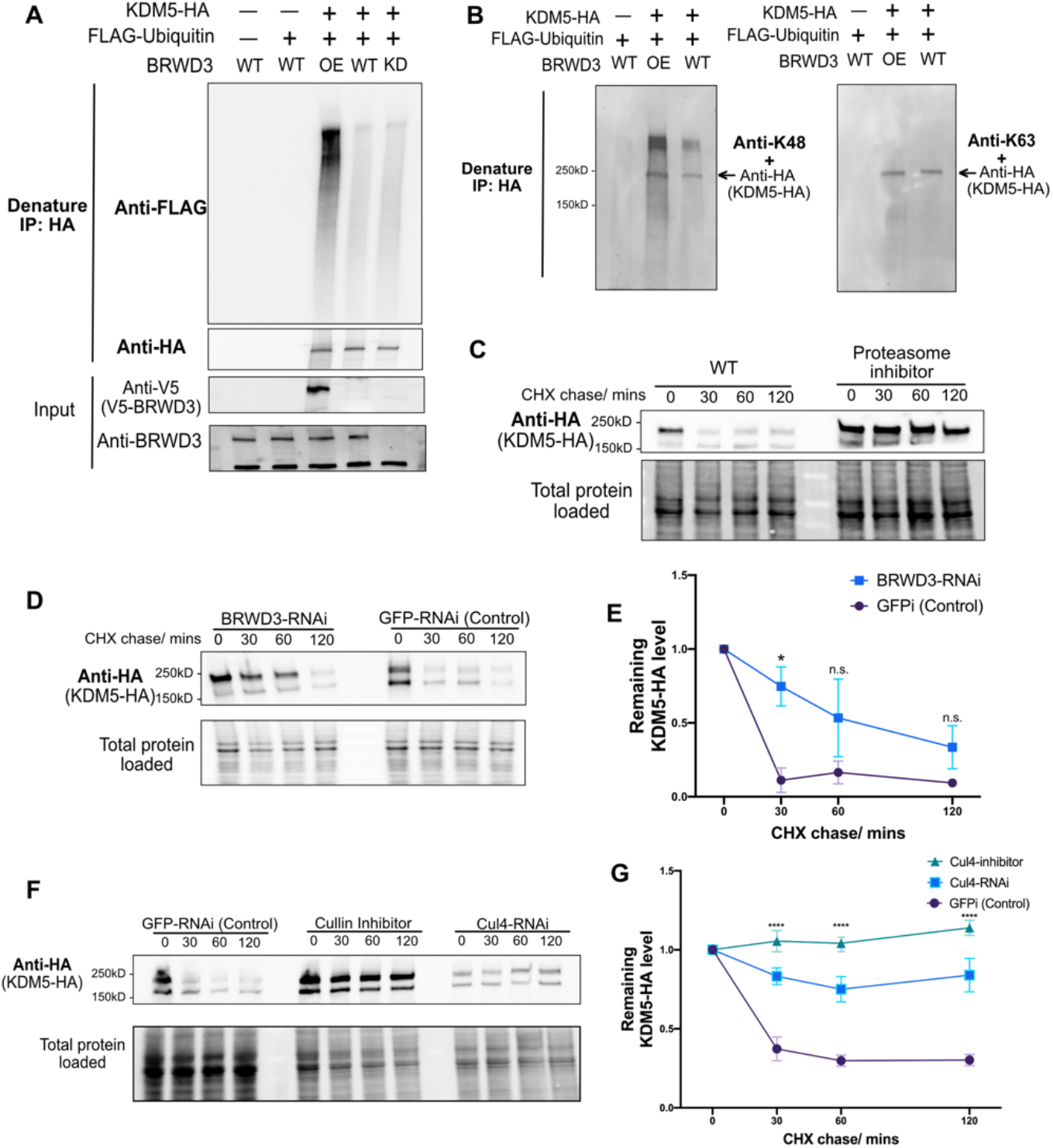
BRWD3 promotes KDM5 ubiquitination and degradation. **(A)** Denaturing IPs from Drosophila S2 cells co-transfected with HA-tagged KDM5 and/or FLAG-tagged ubiquitin in BRWD3-V5 co-transfected (OE), wild-type (WT), or BRWD3 RNAi (KD) conditions. **(B)** Immunoprecipitates from denaturing IPs were blotted with antibodies specific for K48- or K63-linked polyubiquitin chains together with anti-HA antibody. **(C**) S2 cells were transfected with plasmids expressing KDM5-HA and treated with cycloheximide (CHX) only or together with the proteasome inhibitor MG-132. After the indicated time with CHX treatment, Western blots with anti-HA antibody were performed to measure the KDM5-HA level. (**D)** Similar as (C) with either BRWD3-RNAi or GFPi-RNAi treatment. **(E)** Quantification of the KDM5-HA level from three biological replicates of (D) at the indicated time points normalized to the total loaded protein level. **(F)** Similar to (C) in S2 cells with either GFPi- RNAi, Cul4-RNAi, or treatment with the Cullin inhibitor (MLN4924). **(G)** Quantification of the remaining KDM5-HA levels from three biological replicates of (F) at the indicated time points.

To directly test if BRWD3 affects KDM5 stability, we used a cycloheximide chase assay to measure the half-life of KDM5. Cycloheximide inhibits new protein synthesis by blocking translation elongation (43). This allows for the measurement of protein degradation given that new protein synthesis is blocked. Interestingly, we found that KDM5 was rapidly degraded with a half-life of less than 30 minutes (Fig. 3C). Importantly, KDM5 levels could be stabilized by adding the proteasome inhibitor MG-132, demonstrating that KDM5 degradation is proteasome- dependent (Fig. 3C). Next, we tested whether the degradation of KDM5 was dependent on BRWD3. We found that BRWD3 depletion slowed the degradation of KDM5 relative to negative control (Fig. 3D, 3E). KDM5 levels, however, were only partially stabilized upon depletion of BRWD3 and KDM5 was reduced to background levels in ∼120 minutes (Fig. 3D, 2E).

Since we did not find significant change in KDM5 ubiquitination upon BRWD3 depletion, we reasoned that a lesser extent H3K4me2 with little to no activity on H3K4me1 (13, 42). Therefore, we predict that overactive KDM5 removes methyl groups from H3K4me3, ultimately resulting in increased H3K4me1 levels and explaining the altered levels of H3K4 methylation in BRWD3 depleted cells. To test this prediction, we co-depleted KDM5 in BRWD3-depleted S2 cells and measured the relative H3K4me3 by quantitative immunofluorescence. We treated cells with a low dose of KDM5 dsRNA to limit changes in H3K4me3 levels (Fig. S4A). Using a dsRNA concentration that had only a minimal impact on H3K4me3 levels in unperturbed S2 cells, we were able to restore H3K4me3 levels upon BRWD3 depletion (Fig. 4A). To confirm this result, we co-depleted BRWD3 and KDM5 with an independent set of dsRNAs and found H3K4me3 levels were also restored (Fig. S4A). To further test our prediction, we measured the level of H3K4me1 levels in the doubly depleted cells. While we did not observe a full restoration, H3K4me1 levels were significantly reduced upon KDM5 co-depletion (Fig. 4B) (Fig. S4B). BRWD3 is known to associate with chromatin, regulate gene expression and histone modifications (19). Our data indicate that KDM5 is a critical mediator of BRWD3-dependent changes in histone methylation. We wanted to test whether BRWD3 affects chromatin architecture *in vivo* and, if so, if this activity is dependent on KDM5 function. Many genes that regulate chromatin structure and function have been identified as modifiers of position effect variegation (PEV) (45). Interestingly, we found that a *BRWD3* dominantly suppressed PEV. Loss of one copy of *BRWD3* resulted in an increase in variegated *white* expression relative to control flies (Fig. 4C). Thus, we conclude that BRWD3 is a *Su(var)* gene. Critically, loss of a single copy of *KDM5* suppressed the *BRWD3* Su(var) phenotype (Fig. 4C). Consistent with our work in other E3 ubiquitin ligases could also target KDM5 to control its stability (Fig. 3A). To test if KDM5 is degraded in a Cul4- dependent manner, we inhibited Cul4 activity by RNAi or by adding the Cullin-specific inhibitor MLN4924 (44). We found that Cul4 inhibition slowed KDM5 degradation to a greater extent than BRWD3 depletion (Fig. 3F). At 120 minutes, the remaining KDM5 levels in Cul4 depletion were much higher than those in BRWD3 depletion, suggesting that other Cul4 substrate receptors can target KDM5 for ubiquitination (Fig. 3F). Taken together, we conclude that BRWD3 promotes K48-linked polyubiquitin of KDM5 and likely in a Cul4-dependent manner to direct KDM5 degradation.

**Figure 4.**
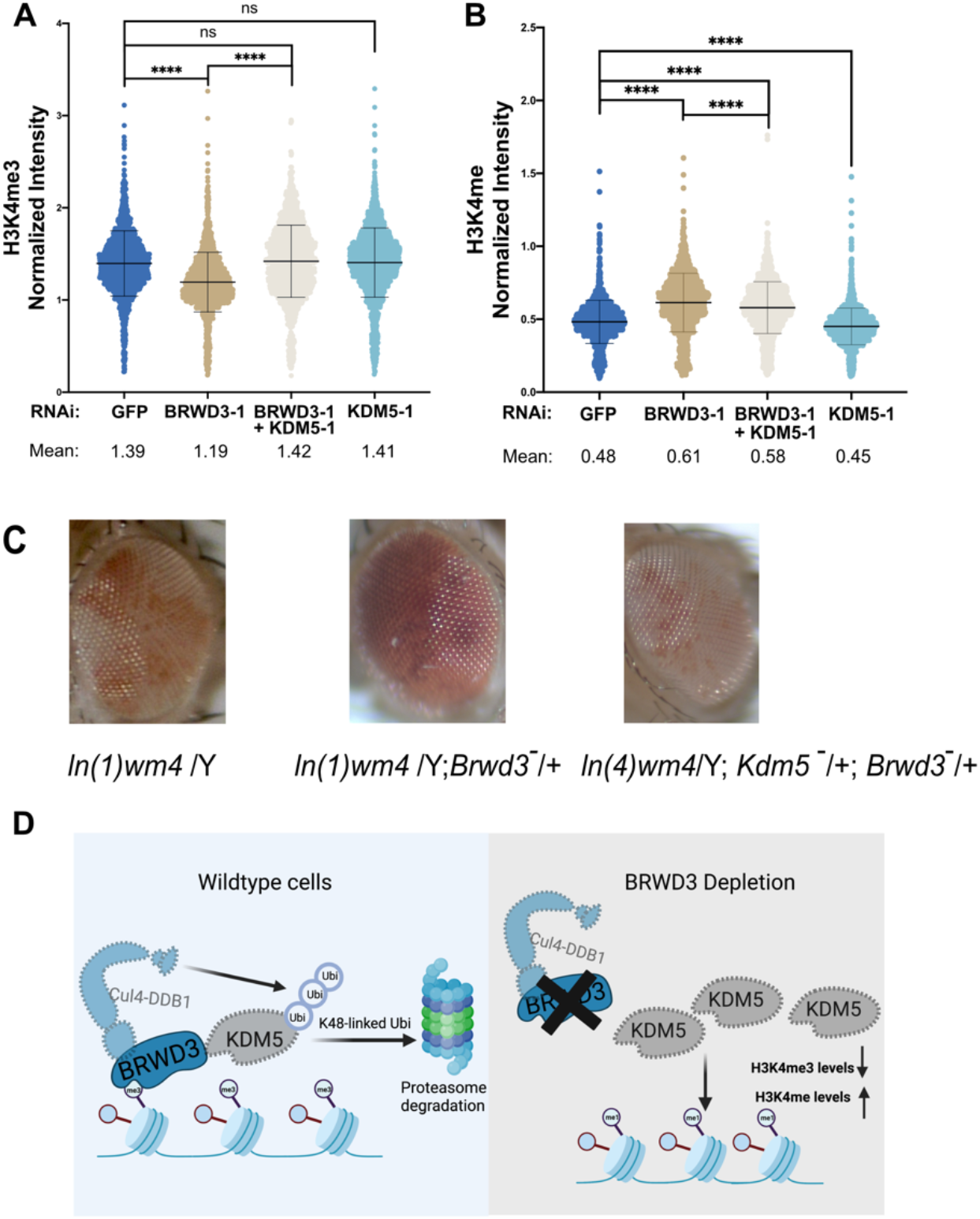
Inhibiting KDM5 restores altered H3K4 methylation levels upon BRWD3 depletion. **(A)** Violin plot of quantitative immunofluorescence assays in Drosophila S2 cells using an anti-H3K4me3 antibody (A) or an anti-H3K4me1 antibody. **(B)** Each distribution represents the signal intensities of 1000 randomly selected cells from three biological replicates. ****p < 0.0001 using the ANOVA one-way analysis with Tukey’s multiple comparisons test. **(C)** Loss of a single copy of BRWD3 gives a Su(var) phenotype that is dependent on KDM5. **(D)** A model for BRWD3-dependent regulation of H3K4 methylation levels. BRWD3 targets KDM5 for degradation. In the absence of BRWD3, KDM5 activity is increased resulting in demethylation of H3K4me3, which generates increased H3K4me1 levels.

### Depleting KDM5 restores altered H3K4me3 levels upon BRWD3 depletion

Given that BRWD3 controls KDM5 stability, we predict that KDM5 is overactive in BRWD3-depleted cells. *In vitro*, KDM5 demethylases preferentially target H3K4me3 and to S2 cells, this indicates that the ability for BRWD3 to affect H3K4 methylation levels and/or function depends on *KDM5*. Taken together, we propose that BRWD3 promotes K48-linked polyubiquitin of KDM5 to promote KDM5 degradation, thus maintaining normal H3K4 methylation levels (Fig. 4D).

## Discussion

Depletion of BRWD3 results in increased H3K4me1 levels (19). The mechanism connecting BRWD3 and H3K4 methylation, however, had yet to be defined. Through several independent approaches, we have found that BRWD3 targets KDM5, a demethylase that targets H3K4me2/me3, for ubiquitination and ultimate proteasomal-dependent degradation. By targeting KDM5, BRWD3 influences not only H3K4me1, but H3K4 di- and tri- methylation. Thus, BRWD3 is a key regulator of pan H3K4 methylation levels. Although we cannot yet conclude that BRWD3 directly targets KDM5 for polyubiquitination, depletion of BRWD3 or Cul4 can stabilize KDM5, indicating that KDM5 polyubiquitination is likely directed by Cul4^BRWD3^. In support of this idea, co-depleting KDM5 upon BRWD3 depletion fully restores changed H3K4me3 levels and partially restores H3K4me1 levels upon BRWD3 depletion. Therefore, we propose that BRWD3 targets KDM5 for degradation to maintain the normal H3K4 methylation levels.

We found that KDM5 is unstable, with a half-life of ∼30mins in Drosophila. We also found that Cul4^BRWD3^ is not the only ubiquitin ligase that targets KDM5 for ubiquitination and ultimate proteasomal-dependent degradation. This suggests that KDM5 activity is highly regulated and KDM5 is likely subject to extraordinary regulation to maintain H3K4 methylation levels. In yeast and human cells, the E3 ubiquitin ligase Not4 has been shown to target KDM5 homologs for polyubiquitination and degradation (48). It is unknown, however, whether Not4 targets KDM5 for ubiquitination in Drosophila. Given that inhibition of Cul4 results in a near complete stabilization of KDM5, it is likely that Cul4 uses other substrate receptors to target KDM5 for polyubiquitination in Drosophila. H3K4me3 is associated with actively transcribed genes and can enhance binding of TFIID to promoters (47, 48). Recently, new evidence suggested that H3K4me3 has role in regulating RNA polymerase II promoter-proximal pause-release (42). H3K4me1, on the other hand, is enriched at enhancers and may serve to recruit chromatin modifying proteins to enable enhancer activity (7, 49). Therefore, carefully regulating the balance of H3K4 methylation states is likely important for gene expression changes. Recently work in mammalian cells has revealed that KDM5 homologs can eliminate nearly all H3K4me3 in less than two hours (42). Given the potential for such rapid demethylation, KDM5 would be an attractive target for extensive post-translational regulation (43). Control of KDM5 activity could be critical for regulating the gene expression programs that occur during cell differentiation. Thus, identifying the additional factors that target KDM5 for degradation will be important to understand how KDM5 activity is regulated during development and disease.

Out of all the histone medications we could assay an unbiased computational approach, we found that H3K4me3 was the most significantly enriched histone modification at BRWD3 binding sites. This is consistent with previous *in vitro* data demonstrating that BRWD3 preferentially binds to H3K4me3 (19, 20). Given that KDM5 is a primary demethylase for H3K4me3/me2, it is possible that BRWD3 binds to H3K4me3 on chromatin and protects H3K4me3 by blocking KDM5 occupancy and targeting KDM5 for degradation. In this case, BRWD3 could form a local zone of protection for H3K4me3, thus increasing the half-life of this mark. In human cells, BRWD3 is necessary to recruit Cul4 onto chromatin. Therefore, it seems likely that the BRWD3- dependent degradation of KDM5 happens in the context of chromatin. Upon BRWD3 depletion, Cul4 would not be properly recruited to chromatin and KDM5 activity would go unchecked, resulting in a ubiquitous decrease in H3K4me3 levels throughout the genome. This is consistent with our findings demonstrating that global levels of H3K4me3 are reduced upon BRWD3 depletion. KDM5 removes methyl groups from H3K4me3 in a sequential manner, first converting H3K4me3 to H3K4me2 and then H3K4me2 to H3K4me1 (13). *In vitro*, KDM5 primarily targets H3K4me3, and to a lesser extent H3K4me2, with limited activity on H3K4me1 (13, 46, 47, 50). Recent *in vivo* time-course assays found that KDM5 enzymes can demethylate H3K4me3 in less than two hours but requires nearly 24 hours to demethylate H3K4me2 or H3K4me1 (42). This supports the finding that KDM5 preferentially targets H3K4me3 *in vitro* (REF). We suspect that the unchecked KDM5 activity upon BRWD3 depletion results in the rapid demethylation of H3K4me3, disrupting the balance of H3K4 methylation and shifting H3K4 methylation towards H3K4me1 at the expense of H3K4me3.

To test our model, we demonstrate that co-depletion of KDM5 can fully suppress the decreased H3K4me3 levels and partially suppress the increased H3K4me1 levels upon BRWD3 depletion. While the H3K4me1 levels were not completely restored upon KDM5 co-depletion, trying to precisely reduce the KDM5 levels with RNAi given the kinetics of H3K4 methylation/demethylation complicate this experiment. Nonetheless, together with all of our data, these results suggest that increased KDM5 activity upon BRWD3 depletion drives changes in pan H3K4 methylation levels. Multiple human genetic studies have demonstrated that human BRWD3 and KDM5C mutations underlie X-linked intellectual disability (17, 18, 29, 30). Further, higher BRWD3 mutation rates have been found in cancers and linked to a decrease in overall survival. Unfortunately, the underlying mechanisms resulting in disease in these cases remains unknown (51, 52). Interestingly, overexpression of KDM5A or KDM5B is also implicated in several cancers progression and drug resistance (16, 53–55). While it is unknown that whether BRWD3 or any of its orthologs promotes KDM5 homolog degradation in humans, it is intriguing to consider that mutations in *BRWD3* could cause increased levels of KDM5. If this is the case, then KDM5 inhibitors could be a potential treatment for *BRWD3*- associated diseases, although further studies will be required to determine whether this pathway is conserved. tristique senectus et netus et malesuada fames ac turpis egestas. Proin semper, ante vitae sollicitudin posuere, metus quam iaculis nibh, vitae scelerisque nunc massa eget pede. Sed velit urna, interdum vel, ultricies vel, faucibus at, quam. Donec elit est, consectetuer eget, consequat quis, tempus quis, wisi.

In in nunc. Class aptent taciti sociosqu ad litora torquent per conubia nostra, per inceptos hymenaeos. Donec ullamcorper fringilla eros. Fusce in sapien eu purus dapibus commodo. Cum sociis natoque penatibus et magnis dis parturient montes, nascetur ridiculus mus. Cras faucibus

## Materials and Methods

### RNA interference

RNAi in S2 cells was performed following previously published methods (56). Briefly, dsRNAs were synthesized using the Invitrogen MEGAscript T7 Transcription Kit (Ambion). A list of primers used to generate dsRNA can be found in Table S2. For each sample, 1.5 million S2 cells were seeded in 1mL non-serum medium in a six-well plate.

20 µg of dsRNA was added and incubated for 45 minutes at room temperature then two mLs of serum-containing medium was added. Cells were incubated for an additional four days. To confirm the efficiency of RNAi depletion, ∼3 million cells were harvested and proteins were extracted by boiling in 2x Laemmli sample buffer (BioRad). Protein levels were measured by Western blotting with different antibodies (Rabbit anti- BRWD3, 1:500).

### Plasmid construction

Plasmid pMT-BRWD3-V5 was created by inserting the BRWD3 CDS sequence into the pMT-puro vector (addgene #17923) using the Gibson assembly method provided by NEB (# E5510S). For the pUbi63E-Flag-Ubiquitin plasmid, a gene block consisting of the pUbi63E promoter and Flag-Ubiquitin CDS sequence was chemically synthesized by Twist Biosciences. Subsequently, the gene block was integrated into the pMT-puro vector by replacing the existing pMT promoter and V5 tag sequences. Finally, the pUbi63E-KDM5-HA plasmid was generated by replacing the Flag-Ubiquitin CDS sequence with the KDM5-HA CDS sequence using Gibson assembly technology provided by NEB (# E5510S).

### Fly stocks and PEV assays

The following lines were acquired from the Bloomington Stock Center: *ln(1)white m4* (BDSC#32578), *BRWD3PX1* (BDSC #35560) and *lid*/*kdm510424* (BDSC # 12367). ORR flies were a gift from Terry Orr-Weaver (Whitehead Institute). Endogenously HA-tagged *BRWD3-HA* fly line was constructed using CRISPR/Cas9-catalyzed homology-directed knock-in method following the online protocol (https://flycrispr.org/scarless-gene-editing/). The 2x HA tag was inserted in the C-terminus of BRWD3 protein. The plasmid injections were performed by BestGene Inc. (https://www.thebestgene.com). For PEV assays, *ln(1)white m4* exhibits variegation of red and white eye color and was used as a control for the experiments. *ln(1)white m4* virgins were crossed to *BRWD3PX1* males and their male offspring were examined for eye color. *ln(1)white m4*/*w-* ; *BRWD3PX1*/*TM3* was crossed to male *kdm510424*/*CyO* and their male offspring were examined for the eye color.

### Quantitative immunofluorescence

RNAi-treated cells were attached to Concanvan A-coated slides for 15 minutes, fixed for 15 minutes in 4% paraformaldehyde and permeabilized for 15 minutes in PBS supplemented with 0.3% Triton-X-100 (PBT). Cells were then blocked for 60 minutes in blocking buffer (1% BSA and 0.2% goat serum in 0.1% PBT). After blocking, cells were incubated with either rabbit anti-H3K4me1 (Cell Signaling #5326, 1:1000), anti-H3K4me2 (Cell Signaling #9725, 1:1000) or anti-H3K4me3 (Active Motif # 39060, 1:1000) antibodies overnight at 4°C in blocking buffer. After washing with PBT, cells were incubated with goat Alexa fluorophore 568-conjugated anti-rabbit IgG secondary antibody (1:500, Life Technologies, # A11011) in blocking buffer for 1h at room temperature. After three washes PBT, cells were stained with DAPI (0.1 µg/mL) in PBT for 10 minutes and mounted in Vectasheild (Vector Labs).

Quantitative immunofluorescence was perform as described previously (57). Briefly, images were obtained using Nikon Ti-E inverted microscope with a Zyla sCMOS digital camera with a 40X oil objective. For each biological replicate, all samples were captured at the same magnification and same exposure time. For quantitative analysis of H3K4 methylation levels, regions of interest (ROIs) were defined based on the DAPI signal. The signal intensity of H3K4 methylation levels were extracted for each ROI. The signal was normalized to the DAPI signal intensity to account for differences in the amount of DNA per cell. For each replicate, three images were randomly taken and quantified. ∼300 randomly selected cells were used for each biological replicate. Two to three biological replicates were used for data analysis. Kruskal–Wallis one-way analysis of variance was performed in GraphPad Prism for statistical significance.

### Immunoprecipitation

Immunoprecipitations (IPs) were performed as previously described (35). Briefly, 1g of flash-frozen embryos aged 18-24h after egg laying (AEL) were collected, dechlorinated and flash frozen in liquid nitrogen. Frozen embryos were ground with a mortar and pestle in liquid nitrogen and then dissolved in 2mL of NP40 lysis buffer (50 mM Tris HCl pH 8.0, 150 mM NaCl, 1% NP40, 1 mM EDTA, 1 mM EGTA) with 2 x Protease Inhibitor Cocktail (Millipore Sigma cOmplete™) on ice. 30 U/ml of benzonase was added and incubated on ice for 30 minutes. The lysate was spin down at 4°C for 5 minutes at 4000 RCF and the supernatant was for IP with 25μL of pre-washed Pierce™ anti-HA magnetic beads (Sigma-Aldrich). After a 2h incubation at 4 °C for 2h, beads were washed three times with NP40 lysis buffer. Proteins were eluted by boiling in 50μL 2x Laemilli sample buffer supplemented with 50 mM DTT for five minutes. Three biological replicates were performed for BRWD3-HA embryos and ORR control embryos. The extracts were labeled with TMT for quantitative mass spectrometry as described below.

### TMT labeling and tandem mass spectrometry

The IP elutes were verified by Western blot (5% of total material) and the remaining material was precipitated using methanol and chloroform. The pellet was washed with methanol to remove excess detergent. Protein was then dissolved in 5μL fresh 1% RapiGest SF Surfactant (Waters Cat. #186001861) and diluted with 10μL 0.5M HEPES pH 8.0 and 32.5μL H2O. 0.5μL of a 0.5M TCEP was added to reduce disulfide bonds. 1μL of a 0.5M iodoacetamide was added to acetylate free sulfhydryl groups for 30min at room temperature in the dark. Trypsin digestion was performed overnight at 37℃ with rigorous shaking. Trypsinized samples were labeled with TMT using a TMT10plex kit (Thermo Scientific catalog #90110). Excess label was neutralized with ammonium bicarbonate at a final concentration of 0.4% for 1h. Samples were mixed and acidified to pH 2 with formic acid. Sample volume was reduced to 1/6th of the original volume using a SpeedVac and restored to the original volume using Buffer A (5% acetonitrile, 0.1% formic acid). Rapigest was cleaved by incubating at 42°C for 1h. Samples were spun at 14,000 rpm for 30 min and supernatant was transferred to a new tube and stored at -80°C.

### In vivo ubiquitination assays

S2 cells were co-transfected with plasmids expressing FLAG-Ubiquitin and HA-tagged KDM5 for two days. For transient transfection in wild-type and *BRWD3* depleted S2 cells, cells were transfected with 100ng of FLAG-Ubiquitin-expressing plasmid and 200ng of HA-KDM5-epxressing plasmid using the Effectene II (Invitrogen) kit according to the manufacturer’s protocol. For BRWD3 overexpression, cells were co-transfected with 200ng/mL of BRWD3-V5-expressing plasmid together with 100ng of a FLAG-Ubiquitin-expressing plasmid and 200ng of an HA-KDM5-epxressing plasmid. To enrich ubiquitinated KDM5 and prevent its degradation, cells were treated with the proteasome inhibitor MG132 for 16 hours before harvesting.

To purify KDM5-HA in denaturing conditions, 2-3 million cells were harvested and lysed with 100μL denaturing buffer (1% SDS, 10 mM Tris HCl, 150 mM NaCl, 1.0% Triton-×100, 1% Sodium Deoxycholate, 0.5 mM EDTA, and 10 mM dithiothreitol) supplemented with 2x protease inhibitor and 30μL/mL Benzonase for 20 minutes on ice, then boiled for 2 minutes. The lysate was centrifuged at 17,000 x g at 4°C for 5 minutes. The supernatant was transferred to a new tube. 900μL NP40 buffer (50 mM Tris HCl pH 8.0, 150 mM NaCl, 1% NP40, 1 mM EDTA, 1 mM EGTA) supplemented with 2x Protease Inhibitor Cocktail. 15μL of anti-HA beads were added to the lysate and incubated at 4°C for 2 hours. The beads were washed in NP40 buffer three times and protein was eluted in 50μL 2x Laemilli sample buffer (with 50 mM DTT) by boiling for 5 minutes. Protein levels were measured by Western blotting with different antibodies (state the antibodies and dilutions).

### Cycloheximide chase assay

First, RNA interference in S2 cells was performed with 20µg of dsRNA targeting *GFP, BRWD3*, or *Cul4* for two days in six-well plates as described above. Cells were then transfected with 200μg/mL *HA-KDM5*-expressing plasmid DNA using the Effectene II (Invitrogen) kit according to the manufacturers protocol. After two days, cells were treated with the translation inhibitor Cycloheximide (Sigma) dissolved in DMSO at 1mg/mL. Cells were harvested at different time points and centrifuged and lysed in 50μL 2 x Laemilli sample buffer (with 50 mM DTT) and boiled for 5 minutes. To inhibit Cullin E3 ubiquitin ligase activity, cells were treated with 50μM of the Nedd8 inhibitor MLN4924 (Selleckchem). To measure KDM5-HA protein levels, 5μL of each sample was used for Western blot analysis with anti-mouse HA (Cell signaling, 1:1000).

### Western blotting

Samples were loaded on 4-15% Mini-protein TGX Satin- free gel (Biorad) for electrophoresis. The gel was activated, imaged and transferred to a low fluorescence PVDF membrane with a Trans-Blot Turbo Transfer System (Biorad). The blot was imaged directly to measure the total loaded proteins. After blocking in 5% fat-free milk in TBST (140 mM NaCl, 2.5 mM KCl, 50 mM Tris HCl pH 7.4, 0.1% Tween 20), blots were incubated with primary antibody for 1h at room temperature. The primary antibodies used are Mouse anti-HA (Cell signaling #2367, 1:1000), Rabbit anti-HA (Abcam ab9110, 1:2000), Rabbit anti K48-linkage Specific Polyubiquitin Antibody (Cell Signaling #4289,1:1000), Rabbit anti K63- linkage Specific Polyubiquitin Antibody (Cell Signaling #5621,1:1000), Mouse ANTI-FLAG® M2 antibody (Sigma Aldrich, 1:1000), Rabbit anti- BRWD3 ((19), 1:500), Rabbit anti-H3K4me3 (Active Motif, 1:1000), Rabbit anti-H3K4me2 (Cell Signaling,1:1000), Rabbit anti-H3K4me1 (Cell Signaling,1:1000). After primary antibody incubation, blots were washed three times in TBST. Blots were then incubated with HRP conjugated secondary antibody (Jackson Labs, 1:15,000) and/or fluorophore conjugated secondary antibodies (Biorad, 1:2,000), washed three times and imaged. To quantify protein levels in Western blot, Bio-Rad Image Lab software was used. Band intensities were measured and normalized to the total amount of protein loaded in each lane (based on the total protein on the blot).

### ChIP-seq analysis

For Figure 1, H3K4me3 ChIP-seq data in wild-type and BRWD3 depleted Drosophila S2 cells (generated in (19); GSE101646) were analyzed. For Figure 1B, the Genome coverage Bigwig files (dm3) were downloaded and visualized using the Integrated Genomics Viewer (IGV) (Broad Institute). For Figure 1C, to acquire H3K4me3 ChIP-seq peaks, H3K4me3 ChIP-seq fastq files in wild-type S2 cells were aligned to dm3 genome using Bowtie2 in Galaxy with default conditions. Duplicate reads were tagged using with Markduplicates (Broad Institute). Peaks were called from alignment result with MACS2 callpeak tool with default settings. Deeptools plotHeatmap was then used to generate the mean ChIP-seq signal plots and heatmaps centered on H3K4me3 peaks (dm3). For Figure 2, ChIP-seq data of BRWD3 (generated in S2 cells by (19); GSE101646), and KDM5 (generated in adult flies by (32); GSE70591) was used. BRWD3 and KDM5 ChIP-seq coverage files were generated using DeepTools BAMCoverage tool with the following options: 1X normalization, bin size = 50 bps, effective genome size = dm6. Genome coverage tracks were visualized using the IGV-Web (Broad Institute). The BRWD3 and KDM5 ChIP-seq peak files were generated using Bowtie2, Markduplicates, and MACS2 tools with the same setting as in Figure 1C except using dm6 genome. The overlap between BRWD3 and KDM5 peaks was generated using the Intersect intervals tools in bedtools. The p-value was calculated with a Fisher test between the two peak files using bedtools FisherBed tools.

### Chromatin marker enrichment in BRWD3 peaks

We downloaded annotations for histone modifications and transcription factor binding sites from modENCODE. The annotations are derived from ChIP-seq and ChIP-chip files in Drosophila melanogaster S2 cells (dm3).

We calculated enrichment for overlap between BRWD3 and modENCODE annotations derived from ChIP-seq/ChIP-chip. Overlap was determined by counting the number of base-pairs overlapping between the BRWD3 peak and the histone modification or transcription factor binding peak using BEDTools (58). We used a permutation-based strategy to determine whether the number of overlapping base pairs was more or less than expected compared to the null distribution.

We generated the null distribution by randomly shuffling the modENCODE annotation peaks throughout the genome. The random peaks were length-matched with the original set of annotated peaks and exclude any blacklisted or gap regions (59). In addition, for modENCODE data obtained from ChIP-chip, we required that the shuffled peaks maintained the probe density of the original peak in addition to matching on the length. We reshuffled peaks that fell more than 0.1 away from the original probe density until at least 99% of the peaks could be appropriately matched. For each BRWD3 peak and modENCODE annotation, we performed 1000 permutations. Finally, we calculated an empirical p value for the observed amount of overlap by comparing it to the null distribution generated above. We correct the p-values for multiple testing using the Benjamini-Hochberg procedure with a false discovery rate < 0.05.

### MudPIT liquid chromatography-tandem mass spectrometry

Triphasic MudPIT columns were prepared as previously described using alternating layers of 1.5cm C18 resin, 1.5cm SCX resin, and 1.5cm C18D (60). Pooled TMT samples (roughly 20 µg of peptide from lysate samples) were loaded onto the microcapillaries using a high-pressure chamber, followed by a 30 minute wash in buffer A (95% water, 5% acetonitrile, 0.1% formic acid). Peptides were fractionated online by liquid chromatography using an Ultimate 3000 nanoLC system and subsequently analyzed using an Exploris480 mass spectrometer (Thermo Fisher). The MudPIT columns were installed on the LC column switching valve and followed by a 20cm fused silica microcapillary column filled with Aqua C18, 3µm, C18 resin (Phenomenex) ending in a laser-pulled tip. Prior to use, columns were washed in the same way as the MudPIT capillaries. MudPIT runs were carried out by 10µL sequential injections of 0, 10, 20, 40, 60, 80, 100 % buffer C (500mM ammonium acetate, 94.9% water, 5% acetonitrile, 0.1% formic acid), followed by a final injection of 90% C, 10% buffer B (99.9% acetonitrile, 0.1% formic acid v/v). Each injection was followed by a 130 min gradient using a flow rate of 500nL/min (0-6 min: 2% buffer B, 8 min: 5% B, 100 min: 35% B, 105min: 65% B, 106-113 min: 85% B, 113-130 min: 2% B). ESI was performed directly from the tip of the microcapillary column using a spray voltage of 2.2 kV, an ion transfer tube temperature of 275°C and a RF Lens of 40%. MS1 spectra were collected using a scan range of 400-1600 m/z, 120k resolution, AGC target of 300%, and automatic maximum injection times. Data-dependent MS2 spectra were obtained using a monoisotopic peak selection mode: peptide, including charge state 2-7, TopSpeed method (3s cycle time), isolation window 0.4 m/z, HCD fragmentation using a normalized collision energy of 32%, resolution 45k, AGC target of 200%, 120 ms (IP) maximum injection times, and a dynamic exclusion (20 ppm window) set to 60s.

### Peptide identification and quantification

Identification and quantification of peptides were performed in Proteome Discoverer 2.4 (Thermo Fisher) using a UniProt *Drosophila melanogaster* proteome database (downloaded February 6th, 2019) containing 21,114 protein entries. The database was adjusted to remove splice-isoforms and redundant proteins and supplemented with common MS contaminants. Searches were conducted with Sequest HT using the following parameters: trypsin cleavage (maximum 2 missed cleavages), minimum peptide length 6 AAs, precursor mass tolerance 20ppm, fragment mass tolerance 0.02 Da, dynamic modifications of Met oxidation (+15.995 Da), protein N-terminal Met loss (−131.040 Da), and protein N-terminal acetylation (+42.011 Da), static modifications of TMT 10plex (+229.163 Da) at Lys and N-termini and Cys carbamidomethylation (+57.021 Da). Peptide IDs were filtered using Percolator with an FDR target of 0.01. Proteins were filtered based on a FDR, and protein groups were created according to a strict parsimony principle. TMT reporter ions were quantified considering unique and razor peptides, excluding peptides with co-isolation interference greater that 25%. Peptide abundances were first normalized based on total peptide amounts in each channel, assuming similar levels of background in the IPs. The abundance-normalized peptide amounts were then normalized to medium peptide amount in each channel. Student t-test was performed between ORR-IP groups and HA-IP groups (p<0.05).

## Supporting information

Han_et_al_supplemental_figures

## Acknowledgements

We thank Bill Tansey and Julie Secombe for critical feedback on this manuscript. This work was supported by National Institutes of Health (NIH) General Medical Sciences awards R35GM133552 to L.P. and R35GM128650 to J.T.N. This paper was typeset with the bioRxiv word template by @Chrelli: www.github.com/chrelli/bioRxiv-word-template

## Author contributions

**DH:** Conceptualization, Methodology, Investigation, Visualization, Formal Analysis, Data Curation, Validation, Supervision, Writing - Original Draft. **SHS:** Investigation, Validation, Formal Analysis. **JPD:** Investigation, Data Curation, Writing - Original Draft. **MLB:** Software, Formal Analysis, Data Curation, Writing - Original Draft. **LP:** Funding acquisition, Supervision, Writing - Review & Editing. **JTN:** Conceptualization, Project administration, Funding acquisition, Supervision, Writing - Review & Editing

## Data Sharing Plan

The mass spectrometry proteomics data have been deposited to the ProteomeXchange Consortium via the PRIDE (61) partner repository with the dataset identifier PXD039708.

