## Supplementary material for "BRWD3 promotes KDM5 degradation to maintain H3K4 methylation levels": Han_et_al_supplemental_figures

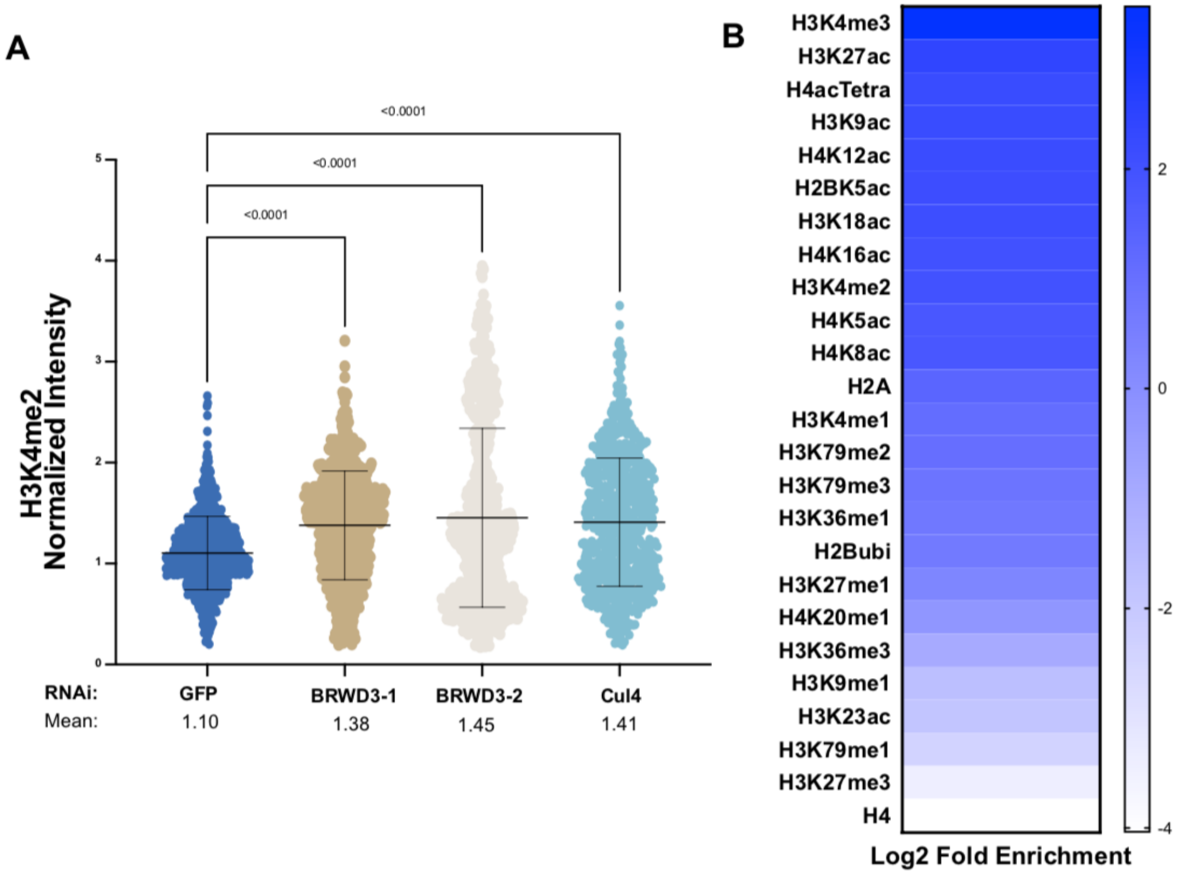


**Supplemental Figure 1. BRWD3 affects H3K4 methylation status.** (A) Violin plots of quantitative immunofluorescence experiments in Drosophila S2 cells using an anti-H3K4me2 antibodies. Each distribution represents the signal intensities of 1000 randomly selected cells from three biological replicates. ****$p < 0.0001$ ANOVA one-way analysis with Tukey's multiple comparisons test. (B) Significant enrichment of all published histone modifications within BRWD3 binding sites. Log2-fold enrichment for observed overlap relative to expected overlap for each mark peak set is shown.

A

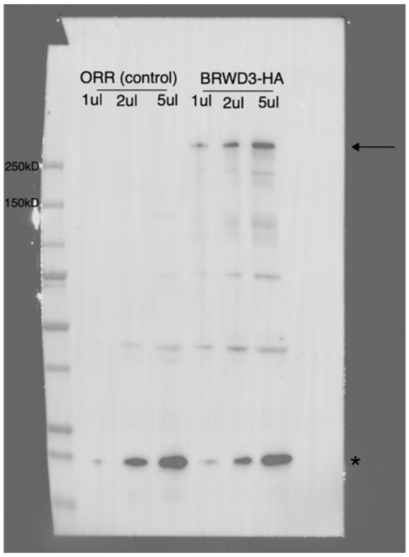


**Supplemental Figure 2. Verification of endogenously HA-tagged BRWD3.** Western blot samples were prepared from endogenously HA-tagged BRWD3 embryos (0-24h) or ORR negative control embryos (0-24h), probing with an anti-HA antibody. Arrow indicates BRW3-HA (~270 kD). Asterisk indicates non-specific bands recognized by anti-HA antibody (both in ORR control and BRWD3-HA).

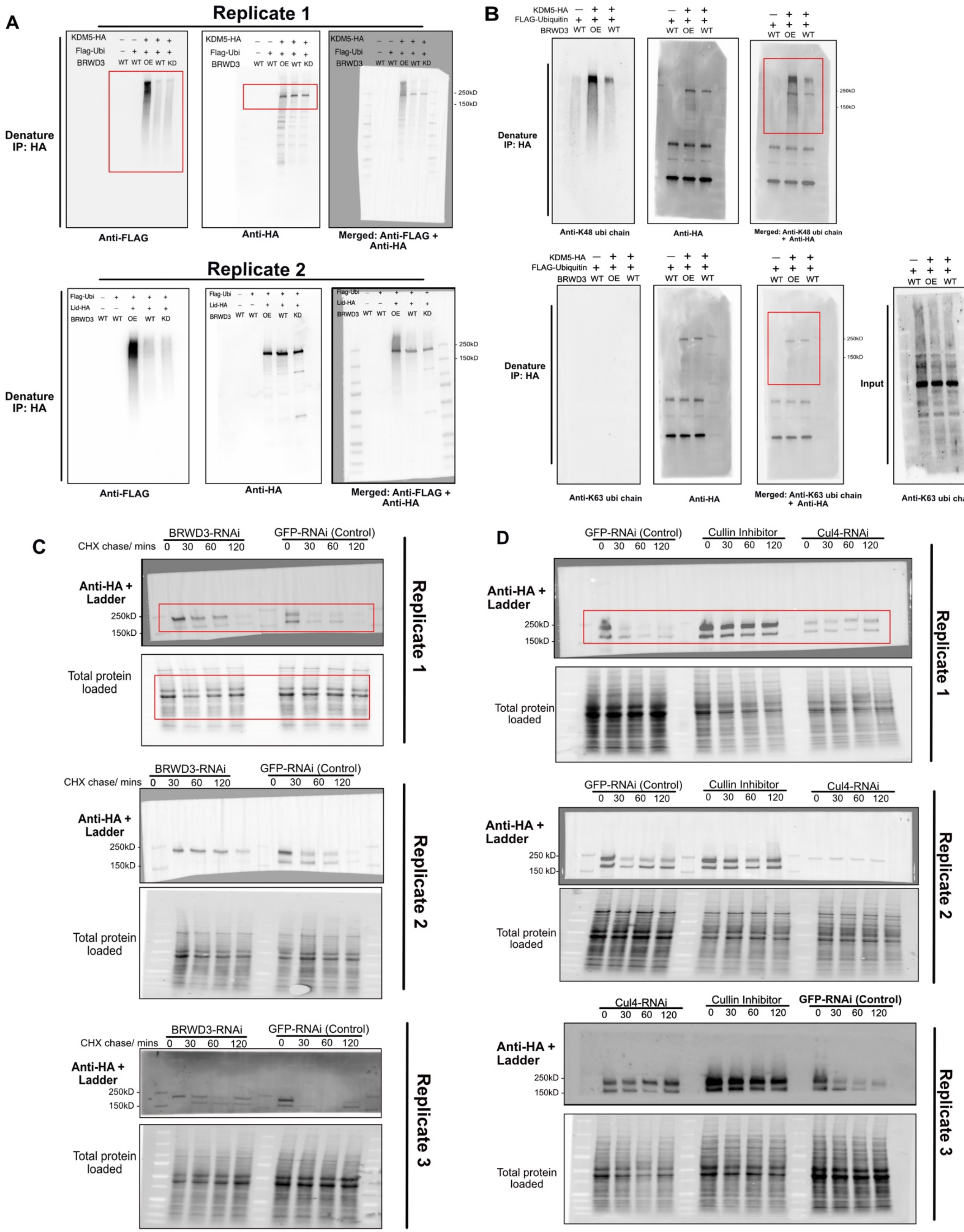
A
Replicate 1
KDM5-HA
Flag-Ubi
BRWD3
WT WT OE WT KD
Denature
IP: HA
250kD
150kD
Anti-FLAG
Anti-HA
Merged: Anti-FLAG + Anti-HA
Replicate 2
Lid-HA
B
FLAG-Ubiquitin
WT OE WT
Anti-K48 ubi chain
Merged: Anti-K48 ubi chain + Anti-HA
Anti-K63 ubi chain
Merged: Anti-K63 ubi chain + Anti-HA
Input
C
BRWD3-RNAi
GFP-RNAi (Control)
CHX chase/ mins
0 30 60 120
Anti-HA + Ladder
Total protein loaded
Replicate 1
Replicate 2
Replicate 3
D
Cullin Inhibitor
Cul4-RNAi

**Supplemental Figure 3. BRWD3 promotes KDM5 ubiquitination and degradation.** Full Western blots and additional replicates used for Figure 3 in the main text. Red boxes indicate cropped regions. (A) Two biological replicates for the denaturing HA-immunoprecipitation followed by Western blots in Fig. 3A are shown. Blots were probed with anti-FLAG (left) and anti-HA (middle) antibodies. The transfection conditions and *BRWD3* expression conditions are indicated on the top. (B) Full Western blots for ubiquitin chain-type specific antibody probing after denaturing HA-IP in Fig. 3B. Separated channels for anti-K48-linked polyubiquitin chains (upper, left) and anti-HA (upper, middle) antibodies, and merged channels (upper, right). Similar order is shown with an anti-K63-linked polyubiquitin antibody (lower). The input samples were blotted with anti-K63-linked polyubiquitin antibody and showed signal (lower, right). (C) Full Western blots for cycloheximide assays under *BRWD3*-RNAi or *GFP*-RNAi conditions used in Fig. 3D. Two more replicates are shown below. (D) Full Western blot for cycloheximide assays under *Cul4*-RNAi, Cullin inhibitor incubation, or *GFP*-RNAi conditions used in Fig. 3F.

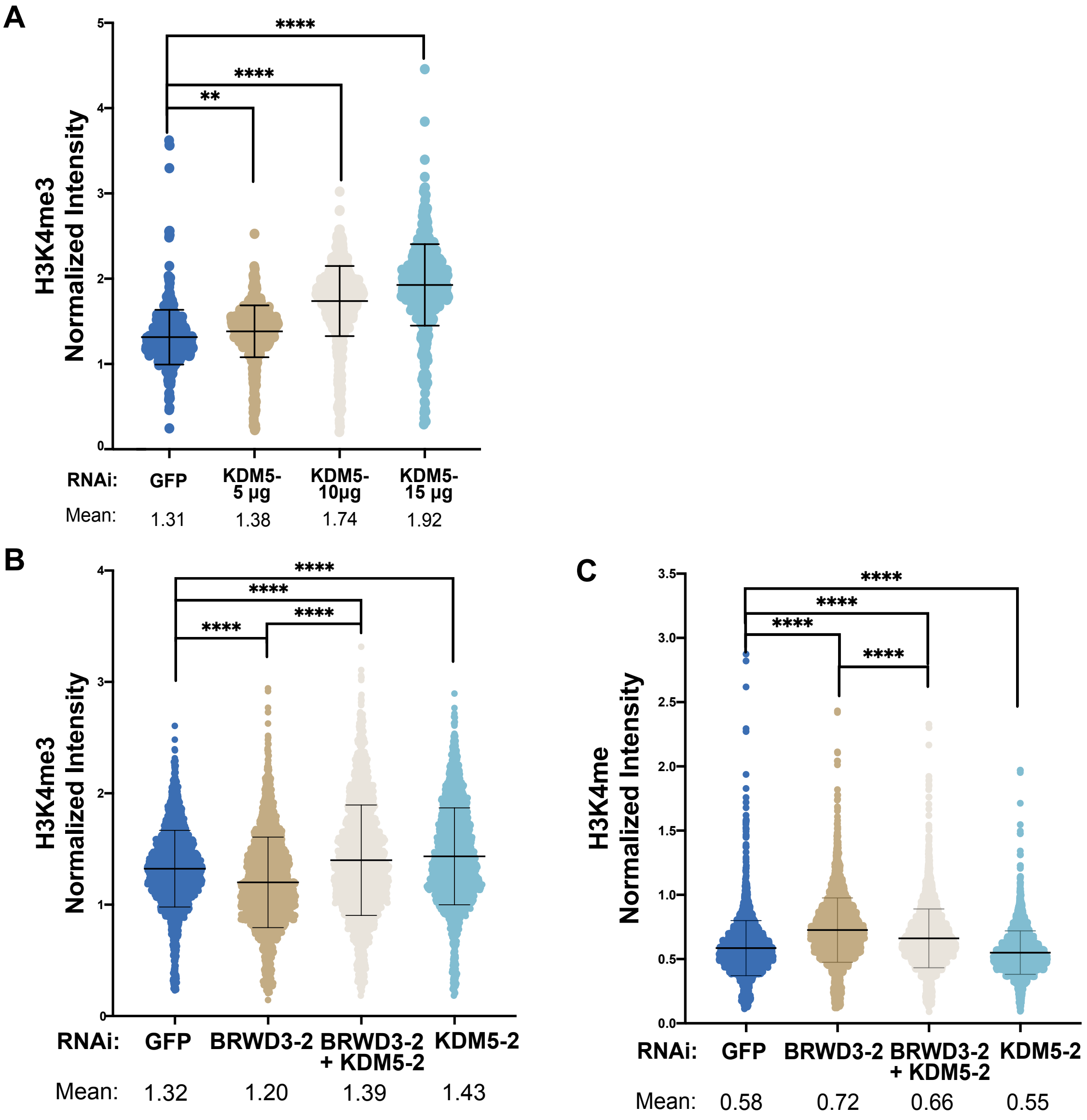


**Supplemental Figure 4. Inhibition of KDM5 with an independent set of dsRNAs restores altered H3K4 methylation levels upon BRWD3 depletion**. Violin plot quantitative immunofluorescence assays plot in Drosophila S2 cells using an anti-H3K4me3 antibody (A, B) or an anti-H3K4me1 antibody (C) with different dsRNA treatments. **p< 0.01, ****p < 0.0001 using ANOVA one-way analysis with Tukey's multiple comparisons test.
